# Barcoded-Plasmid DNA library construction for recording cell lineage trees enabled by a Scalable and modular Biofoundry-based Automated Robotic Pipeline

**DOI:** 10.64898/2026.07.07.736956

**Authors:** Eleonora Tassinari, Lesley Ives, Erica Hawkins, Davide Annese, Sonia Fonseca, Yuxuan Lan, Wilfried Haerty, Edyta E. Wojtowicz, Carolina Grandellis

## Abstract

High-quality plasmid DNA purification at high throughput remains a significant bottleneck in molecular biology and bioengineering. Current methods frequently fail to deliver sufficient yields of pure, transfection-grade DNA required for genetic engineering applications in mammalian cells. Here, we present a Biofoundry-based automated pipeline using the CyBio FeliX robotic liquid handling platform to rapidly purify plasmid DNA with minimal manual intervention. The protocol leverages Solid Phase Reversible Immobilisation (SPRI)-based magnetic bead technology to ensure consistency, scalability, and DNA purity suitable for downstream viral particle production and mammalian cell transfection. The pipeline supports flexible processing of between 8 and 96 samples per run, making it adaptable across a wide range of experimental scales. The protocol is openly available via Earlham Institute GitHub repository, enabling broad adoption across the bioscientific community and contributing to the growing toolkit of reproducible, scalable engineering biology workflows. In this work, we employed an integrated robotic pipeline to process 528 pooled DNA plasmids and built a Lentiviral DNA plasmid library for lineage tracing, validated the library by sequencing, and demonstrated efficacy in downstream mammalian cell transfection experiments.

## Introduction

Plasmid DNA, naturally occurring in bacteria and some eukaryotes, is broadly used in cell biology as a convenient and compact carrier for modifying the genetic makeup of cells. Since their discovery, plasmid vectors have revolutionised molecular biology, enabling researchers to probe the molecular basis of life and underpinning countless advances across basic and applied sciences^1^. Plasmids can be engineered for diverse applications, including gene expression studies, synthesis of recombinant proteins, genetic screens, and gene editing. Particularly in synthetic biology, where high-throughput (HT) preparation is crucial to enable parallel testing of multiple variants, plasmid DNA isolation at scale is essential, as low throughput restricts the pace of design-build-test (DBT) cycles and reduces the number of designs that can be explored in parallel^2^. Applications requiring higher concentrations (typically ≥1–2 µg/µL) of transfection-grade DNA present challenges for HT pipelines due to the typically lower quality of plasmid DNA obtained at scale ^3^. Current HT plasmid preparation methods meet the minimal requirements (20-100 ng/µl) for applications such as Nanopore or Sanger sequencing and DNA assembly ^4^, but more demanding applications, such as mammalian cell line transfection and cell-free protein synthesis (CFPS) ; require higher DNA concentration (200 ng/µl) yield and purity (A260/A280 Ratio 1.8–2.0 and A260/A230 Ratio 2.0–2.2) which remain difficult to achieve at HT scale ^5^. Addressing these limitations requires a more integrated approach to automation and workflow design, an area where Biofoundries are uniquely positioned to advance the field.

Biofoundries are integrated facilities that combine molecular biology and automation, serving as hubs for research and technology development by accelerating biological sample processing at scale across tens of thousands of samples ^6,7,8,9^. The implementation of robotics provides accuracy and precision, supporting the engineering biology field and accelerating the iterative Design-Build-Test-Learn (DBTL) cycle by developing methods, standards, and pipelines to perform those cycles efficiently across laboratories.

While many Biofoundries share similar automated equipment, they often house a unique set of instruments curated based on their specific processes, applications, and areas of expertise. An essential capability across Biofoundries is the efficient extraction of high-quality genomic and plasmid DNA in a high-throughput manner to support downstream processes. While broadly automated workflows for genomic DNA extraction for NGS are available and adapted to various robotic platforms, such as the Opentrons Flex High-Throughput NGS Workstation, the QIAcube Connect, and the Tecan DreamPrep® NGS; equivalent automated protocols for plasmid DNA purification are less common and are typically limited to smaller scales and manual processing.

To support downstream applications including mammalian cell transfection, there is an unmet need for scalable protocols that yield high-quality, concentrated plasmid DNA at scale.

Several automated platforms for high-throughput plasmid DNA purification exist, utilising a range of commercial and custom extraction kits ^10^. However, these approaches face key limitations: sample throughput is often restricted, for example, the QIAcube Connect processes a maximum of 12 samples per run and the PhyNexus AutoPlasmid MEA up to 36 ^10^. More scalable systems, such as the Automated Miniprep Plasmid Station (AMPS) developed by Cohen et al., are tied to specific liquid handling platforms^5^. A further challenge is the heterogeneity of liquid handling equipment across Biofoundries, which hinders protocol transferability ^11^. Collectively, these limitations highlight the need for a scalable, flexible automated plasmid purification pipeline suitable for high-throughput applications.

The CyBio FeliX was identified as the most suitable platform within our Biofoundry for automated plasmid DNA purification, capable of processing between 8 and 96 samples per run. The CosMCPrep kit (Beckman Coulter), based on Solid Phase Reversible Immobilisation (SPRI) technology, was selected for its compatibility with robotic implementation and its proven performance in nucleic acid extraction, purification, and size selection.

In this study, we present an optimised automated workflow using the CyBio FeliX robotic platform for the purification of plasmid DNA from up to 96 samples using the CosMCPrep kit in under 3 hours with minimal manual input. The protocol is openly available via Earlham Institute GitHub repository (https://github.com/EarlhamInst/Plasmid_Preparation_Using_CyBioFelix).

The purified DNA was shown to be of sufficient quality for downstream mammalian cell transfection following a single pooled clean-up step. We anticipate that this workflow will be broadly adopted by the synthetic biology community as an accessible, scalable, and reproducible solution for high-throughput plasmid DNA purification. As plasmid characteristics such as copy number, sequence identity and size may influence extraction efficiency, a proof-of-concept validation is advisable to new users when utilising this pipeline. We anticipate that this workflow will be broadly adopted by the synthetic biology community as an accessible, scalable, and reproducible solution for high-throughput plasmid DNA purification.

## Results

### Design and Construction of a Golden Gate-Compatible Lentiviral Barcoded Backbone Vector

In order to design a cloning strategy compatible with automation pipelines, a modified version of the lentiviral vector pEGZ2^12,13^ was generated to be Golden Gate-compatible. The vector was domesticated and equipped with a unique DNA barcode. An mScarlet dropout cassette was integrated downstream of the Woodchuck Hepatitis Virus Post-Transcriptional Regulatory Element (WPRE) of the pEGZ2 vector to facilitate modular cloning. The mScarlet gene encodes a monomeric red fluorescent protein (RFP), exhibiting excitation and emission maxima of 569 nm and 594 nm, respectively, and is widely used as a fluorescent reporter in mammalian cell applications due to its high brightness, fast maturation, and low cytotoxicity. Using this backbone, additional modifications (see Methods) were introduced through Golden Gate assembly (Figure 1), and the resulting construct was designated EB04119 (https://doi.org/10.5281/zenodo.20719824) and verified by sequencing.

**Figure 1.**
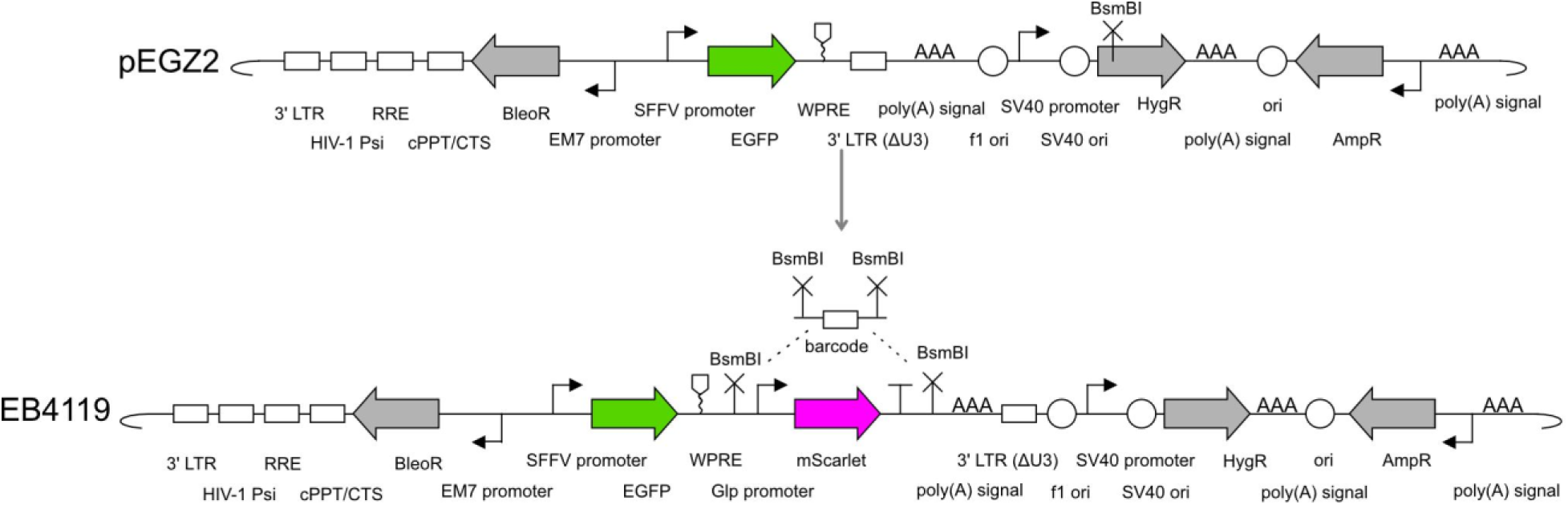
Schematic of lentiviral vector constructs EB04119 and pEGZ2. EB04119 contains a Spleen Focus-Forming Virus (SFFV)-driven EGFP cassette, a bacterial GlpT promoter driving expression of the glycerol-3-phosphate transporter, a mScarlet dropout cassette flanked by BsmBI sites for barcode insertion via Golden Gate assembly, and an SV40-driven hygromycin resistance gene (HygR), within a lentiviral backbone comprising the 3’ LTR, HIV-1 Psi, RRE, cPPT/CTS, WPRE, and ampicillin resistance gene (AmpR). The parental vector pEGZ2 (upper panel) was domesticated for BsmBI restriction enzyme and shares the same backbone but lacks the mScarlet cassette. The red arrow indicates the site of modification. Schematic representations follow the Synthetic Biology Open Language (SBOL) visual standard.

### Development of an Automated Pipeline for Plasmid DNA Purification in 96-Well Format

An automated script for plasmid DNA purification was written and validated for the CyBio FeliX robotic platform. The CosMCPrep Plasmid Prep kit (Beckman Coulter) was selected as it is based on SPRI purification and is therefore well suited to automation. A schematic overview of the automated plasmid DNA extraction pipeline is described in Figure 2. The pipeline input is a 96-deep well source plate containing bacterial pellets and it yields a 96-well output plate containing purified eluted DNA (Figure 2A). The DNA extraction steps comprise the sequential addition of lysis and neutralisation buffers to bacterial pellets, a centrifugation step, automated supernatant transfer, magnetic bead-based binding, ethanol washes, and final elution (Figure 2B). The initial robot deck configuration (Figure S1) incorporates the heater-shaker and AlpAqua magnetic adapter modules. During protocol optimisation, we observed that liquid dripping from the tips caused contamination and potential damage to the platform surface. To address this, disposable absorbent racks were positioned beneath the tip racks and secured to the deck base with adhesive tape, which effectively prevented tip dripping and ensured stable placement of the tip racks throughout the run. Checkpoints were incorporated throughout the protocol as pauses to ensure that (1) bacterial pellets are fully dissolved in buffer RE1, and (2) residual ethanol has evaporated sufficiently prior to elution. These checkpoints can be removed using the CybioComposer software if a more walkaway-compatible process is desired.

**Figure 2.**
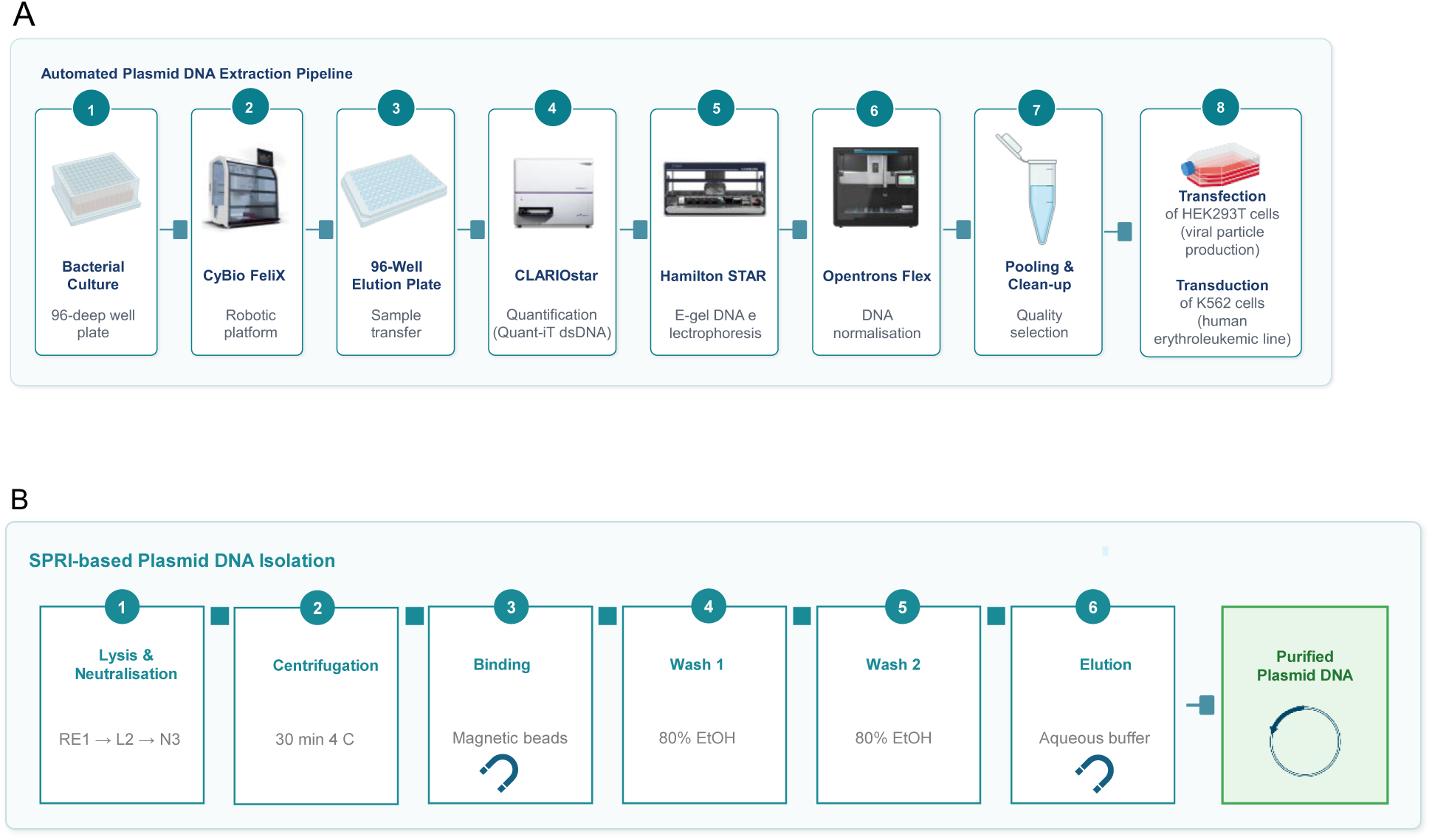
(A) Schematic overview of the automated plasmid DNA extraction pipeline. Bacterial cultures grown in 96- deep well plates are processed using the CyBio FeliX robot, transferring samples into 96-well destination plates. DNA samples were then quantified using the Quant-iT™ dsDNA assay and fluorescence was measured on a CLARIOstar® microplate reader (BMG Labtech). Hamilton and Opentrons Flex systems are used for DNA normalisation and automated electrophoresis. Purified plasmid DNA is pooled, cleaned up and prepared for downstream applications, including transfection of HEK293T cells for viral particle production and transduction of the K562 human erythroleukemic cell line. (B) Schematic representation of the individual SPRI-based DNA isolation steps. Bacterial pellets are resuspended and lysed sequentially using reagents RE1, L2, and N3 (1). Following lysis, a centrifugation step is performed (2). Magnetic bead-based binding selectively captures plasmid DNA (3), which is retained during two successive wash steps to remove contaminants (4 and 5). Purified plasmid DNA is eluted in aqueous buffer, yielding high-quality DNA ready for downstream use (6).

### Automated Extraction of Transfection-Grade Plasmid DNA

The protocol was first tested on 96 plasmid DNA samples from a lentiviral barcoded library constructed using the vector EB04119, which contains a carbenicillin resistance cassette, an EGFP reporter, and a 70-nucleotide barcode for downstream lineage tracing of bone marrow cells in mice. A full 96-well plate of lentiviral vectors was processed for plasmid extraction and quantified using both NanoDrop and Quant-iT™ fluorometric assays. Concentrations determined by Quant-iT™ ranged from 8.8 to 127.0 ng/µL. Of the 96 extractions, 3 (3.2%) failed, 3 (3.2%) yielded very low concentrations (< 20 ng/µL), 79 (82.3%) yielded medium concentrations (20–100 ng/µL), and 11 (11.5%) yielded high concentrations (> 100 ng/µL), with an overall mean concentration of 67.53 ng/µL and a median of 68.9 ng/µL. Although the mean concentration was higher with the manual method (80.28 ng/µL), the median concentration was similar (69.8 ng/µL) suggesting comparable typical yield. Manual extraction showed a higher failure rate (6.3%); among successful extractions, 2 samples (2.1%) had low concentrations, 67 (69.8%) medium concentrations, and 21 (21.9%) high concentrations (percentages may not total 100% due to rounding). Automated extraction was more consistent, with a lower coefficient of variation than the manual protocol (37.4% vs 47.9%) (Figure 3A).

**Figure 3.**
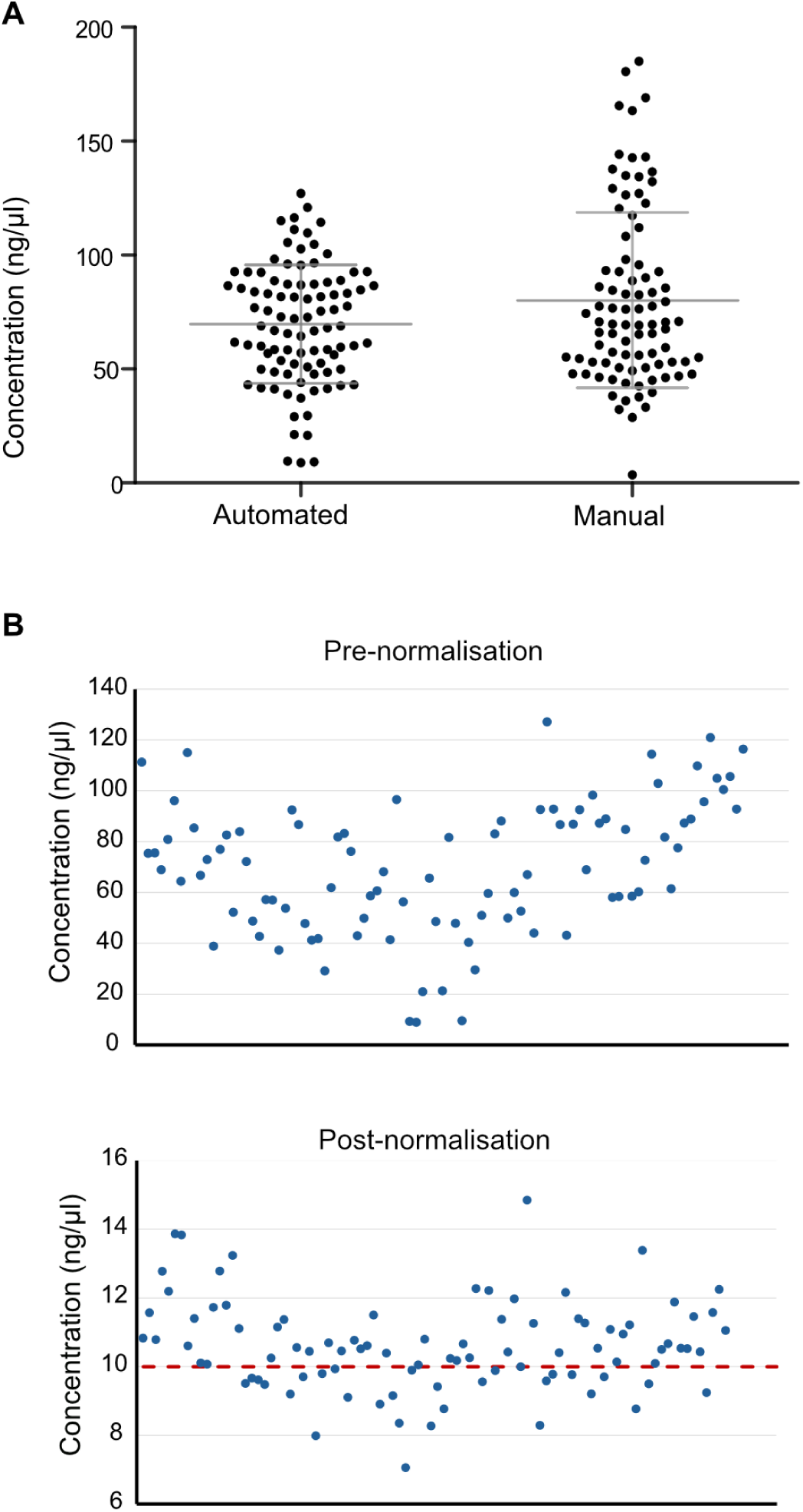
(A) Distribution of the DNA yields (ng/µl) obtained with the automated and manual extraction performed with the CosMCPrep kit (Beckman Coulter). The central horizontal bar represents the average concentration, while the whiskers the standard deviation. (B) Distribution of measured DNA concentrations across samples before and after normalisation to a target concentration of 10 ng/µL (red dashed line) using the Opentrons Flex.

Samples were subsequently normalised to 10 ng/µL using the Opentrons Flex liquid handling platform; and quantified before and after normalisation (Figure 3B). DNA integrity and expected fragment sizes were confirmed by agarose gel electrophoresis using E-Gel™ 96 Agarose Gels (Figure S3). A total of 528 plasmid DNA extractions (5.5 96 well-plates) were successfully completed. DNA was quantified and normalised, with quality assessment confirming that samples met the required standards for downstream applications. All plates yielded consistent results; data from one representative plate are presented here as an example since full library quality control and comprehensive sequencing analysis are beyond the scope of the present study. Following quality control, all wells were pooled into a single tube using a multichannel pipette and subsequently cleaned up using DNA column purification, yielding a final DNA concentration of 109.1 ng/µL (A260/280 = 1.94; A260/230 = 2.18) in a total volume of 15 µL, corresponding to a total yield of 1.64 µg. Pooled and further purified samples were used for transfection of HEK293T cells, achieving successful viral particle production and transduction of the K562 human erythroleukaemic cell line. Successful transduction was confirmed by flow cytometry (FACS), which detected robust eGFP expression, indicating high transduction efficiency (Figure 4).

**Figure 4.**
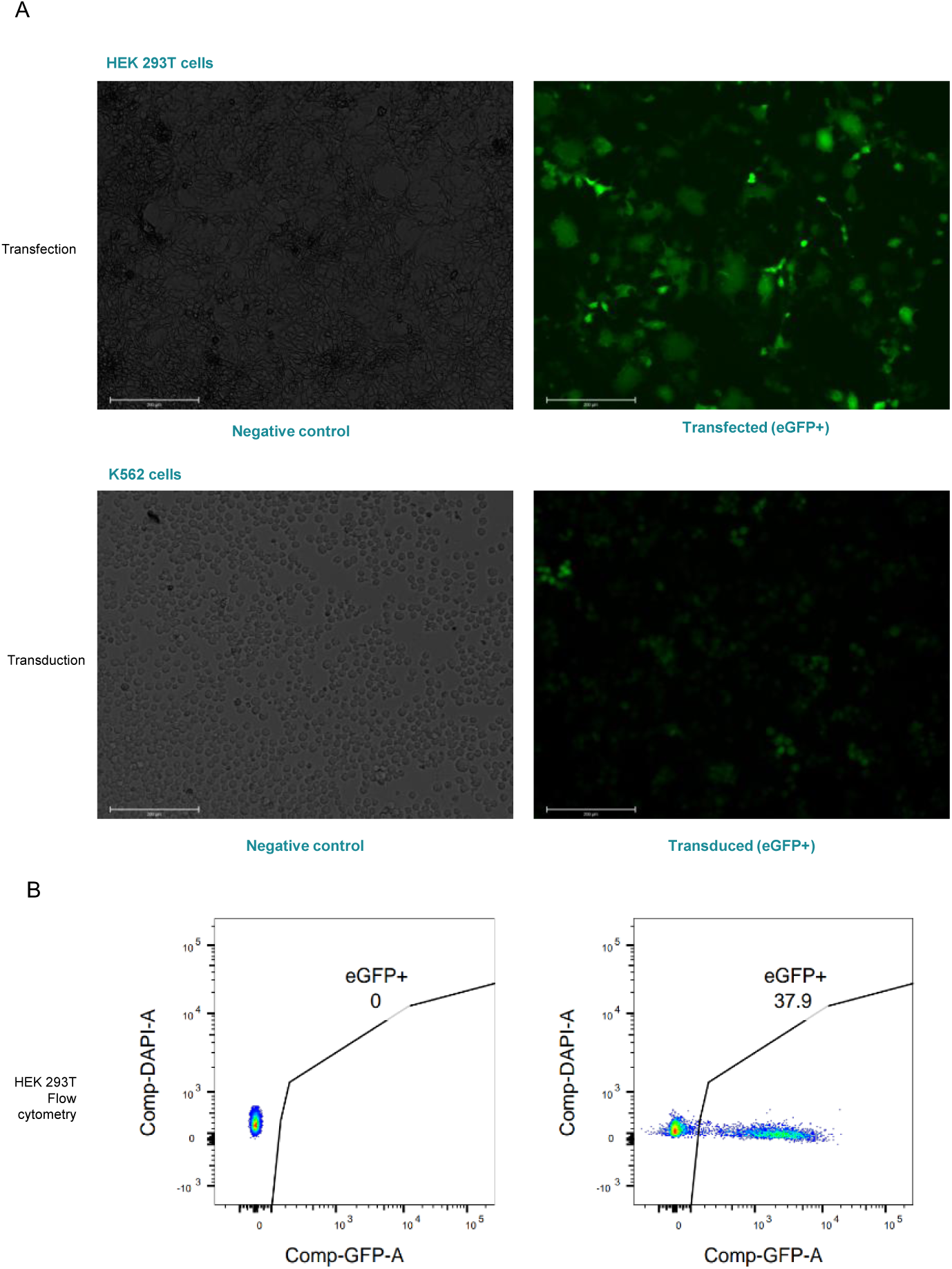
Validation of lentiviral barcoded vector libraries via transfection and transduction into mammalian cells. Fluorescence microscopy images showing GFP expression following (A) transfection of HEK293T cells with barcoded lentiviral plasmid libraries and transduction of K562 cells with the resulting viral particles. Successful GFP expression confirms efficient delivery of barcoded constructs into target cells. Images are representative fields of view; scale bars represent 200 µm (A) and 200 µm (B). Negative control microscopy images are included, showing cell lines that were neither transfected nor transduced. (B) Transduction efficiency was further quantified by flow cytometry (FACS) analysis of eGFP-positive cells. FACS dot plots showing the percentage of eGFP+ K562 cells in the singlet/viable gate.

### High-Throughput Validation of a Barcoded Plasmid Library via Illumina Amplicon Sequencing and bioinformatic validation

To confirm the completeness of the barcoded plasmid library, we used a high-throughput quality control and validation pipeline. This consisted of high-throughput PCR amplification across all plasmid pools, quantitative assessment by qPCR, and amplicon sequencing by Illumina, followed by bioinformatic analysis to verify barcode representation across the library. Sequencing analysis yielded over 97% of paired end reads successfully merged into single reads. Following barcode extraction and filtering, barcodes were validated against the expected library structure, with low-frequency barcodes and low-quality wells removed to ensure data integrity (Table S2). In total, 2,255 barcoded plasmids were validated. To functionally validate the library HEK293T cells were transfected with the barcoded library (Figure 4A), the obtained virus population was used to transduce K562 cells (Figure_4A-B). Successful eGFP fluorescence detection following transfection and transduction of cell lines further confirmed library functionality. To further validate the barcode composition in transduced K562 cells, an aliquot of cells was sequenced. Following barcode extraction and filtering 2,255 barcodes were detected (Table S3), demonstrating that the pipeline constitutes a robust and novel automated tool for lineage tracing barcoded DNA library preparation.

## Discussion

In this study, we present a scalable, modular, and automated pipeline for high-throughput plasmid DNA purification using the CyBio FeliX robotic liquid handling platform, that was used to process 528 DNA plasmid quadruples and built a Lentiviral DNA plasmid library for lineage tracing. The library was validated by sequencing, and we demonstrated its efficacy in downstream mammalian cell transfection and transduction experiments by fluorescence activated cell imaging and by Illumina sequencing with a total number of 2,255 barcodes identified. To our knowledge, this represents one of the few openly available automated plasmid purification workflows for library preparation based on Solid Phase Reversible Immobilisation (SPRI) technology capable of processing up to 96 samples in a single run with limited manual intervention. A key advantage of SPRI-based magnetic bead purification over conventional silica column-based approaches is its flexibility: unlike filter plate-based methods, which require multiple centrifugation steps and are constrained to processing full 96-well plates, the magnetic bead-based workflow supports processing of between 8 and 96 samples per run^14^ contributing to more sustainable laboratory practices^11^. This modularity is valuable in Biofoundry settings, where sample numbers and experimental requirements vary between projects. In addition, silica column-based alternatives are less flexible and sustainable, generating greater plastic waste and lacking the scalability required for Biofoundry applications. We then selected the CosMCPrep kit (Beckman Coulter) as the most suitable robotic platform. The automated SPRI-based pipeline developed here addresses several key limitations identified in previously reported approaches.

Cohen et al. developed the Automated Miniprep Plasmid Station (AMPS), which produced high-quality, high-yield DNA suitable for downstream mammalian expression and antibody discovery screening, significantly accelerating the DBTL cycle^5^. However, this pipeline was developed specifically for the Biomek i7 hybrid liquid handler and could not be adapted to our facility. The QIAcube Connect and PhyNexus AutoPlasmid MEA ^10^ are further constrained by limited throughput of 12 and 36 samples per run, respectively, whereas our pipeline processes up to 96 samples in a single 3-hour run, representing a substantial improvement in scalability and accessibility. Suckling et al. developed a high-throughput plasmid library preparation method using the Labcyte Echo® system and Design of Experiments optimisation ^15^; however, this approach requires six centrifugation steps, limiting walkaway time, increasing the staff time and associated cost. Our pipeline requires only a single manual centrifugation step, which minimally interrupts the automated walkaway workflow and does not constitute a critical limitation, as the speed and timing parameters are fixed. Our protocol was developed on the CyBio FeliX, which is a compact, accessible mid-throughput platform; and the pipeline is openly available via GitHub to facilitate adoption across laboratories. We acknowledge that translating complex automated protocols between liquid handling platforms remains a significant barrier to the decentralisation of synthetic biology workflows ^16^, and we anticipate that open availability of protocols and the use of mid-range affordable automation platforms will help address this challenge. In the future, full integration of centrifugation into the automated workflow could be explored, although this would likely require different instrumentation. The overall success rate of the pipeline was high, with 96.8% of extractions yielding detectable DNA. Of these, 82.3% fell within a medium yield range (20–100 ng/µL) and 11.5% exceeded 100 ng/µL, with an overall mean concentration of 67.53 ng/µL as measured by Quant-iT™ fluorometric quantification.

Only 3.2% of extractions failed entirely, and a further 3.2% yielded very low concentrations (< 20 ng/µL). These failure rates are consistent with those reported in comparable automated pipelines and are likely attributable to variability in bacterial culture density or pellet quality rather than to limitations of the automated protocol itself. The inclusion of operator checkpoints at critical steps, specifically to confirm complete pellet resuspension in buffer RE1 and sufficient ethanol evaporation prior to elution; helped reduce protocol failures. These checkpoints can be removed to suit laboratories at different stages of automation maturity, from supervised to fully unattended operation. Downstream validation confirmed that the DNA produced by this pipeline is of sufficient quality for transfection-grade applications following normalisation to 10 ng/µL using the Opentrons Flex, pooling, and a single clean-up step. Although this additional clean-up is required prior to transfection, it is performed only once on the pooled sample rather than on individual wells, representing a straightforward and rapid manual step that has negligible impact on overall walkway time. The purified pooled DNA achieved a concentration of 109.1 ng/µL with A260/280 and A260/230 ratios of 1.94 and 2.18, respectively, indicating high purity and low protein or solvent contamination. Successful transfection of HEK293T cells and subsequent transduction of K562 cells, confirmed by eGFP expression detected by fluorescence microscopy and flow cytometry (FACS), demonstrate that the pipeline reliably produces DNA suitable for downstream mammalian cell applications. High DNA quality was confirmed by sequencing, and the 2,225 barcodes detected indicate well-balanced barcode representation within the generated library. The elevated barcode count observed in K562 samples likely reflects insufficient sequencing depth for this sample: its barcode complexity was 50-fold higher than all other samples, yet the sequencing yield did not scale accordingly. This can be addressed in future experiments by allocating greater sequencing depth to samples transduced with the full library.

The generated library represents an advance over many existing high-throughput plasmid preparation methods, which typically yield DNA suitable for sequencing or DNA assembly but insufficient for mammalian cell transfection. The integration of the Hamilton STAR liquid handling system for automated gel loading further streamlined quality control, enabling simultaneous assessment of DNA integrity across all 96 samples in a single step. Combined with fluorometric quantification and automated normalisation on the Opentrons Flex, this multi-instrument approach exemplifies the modular nature of the pipeline, in which individual steps can be adapted or replaced depending on the equipment available at a given Biofoundry. This is particularly relevant given the well-documented challenge of protocol transferability across biofoundries ^11,16^, arising from the considerable heterogeneity of liquid handling platforms available commercially. Looking ahead, several improvements could further enhance the pipeline. Incorporation of fully automated centrifugation steps would reduce remaining manual handling and increase true walkaway time. Optimisation of elution volumes and bead ratios for specific plasmid sizes or backbone types could improve yields for specific applications however this is plasmid dependent and needs further validation. The automated pipeline described here represents a valuable addition to the Biofoundry toolkit, enabling reliable, scalable, and high-quality plasmid DNA purification that supports the DBTL cycle and broadens access to transfection-grade library DNA preparation. Biofoundries have a unique opportunity to harness the power of automated biological workflows, and the establishment of the Global Biofoundry Alliance has created an ecosystem for open technologies to be developed and shared, facilitating the generation of common standards, open-source reference materials, and tools across the global community^7,11^. Importantly, while validated here in the context of mammalian cell applications, biological pipelines inherently require re-validation when applied to different systems or host contexts, such as plant or other mammalian systems. By making the protocol openly available through our GitHub repository (https://github.com/EarlhamInst), ee aim to encourage widespread adoption across the bioscience community. We envisage that this workflow could serve as a foundation for integrating plasmid DNA preparation into larger, multi-step automated DBTL.

## Acknowledgements

This work was funded by the BBSRC grants BB/E/ER/23NB0007 (BBSRC National Bioscience Research Infrastructure: Earlham Biofoundry) awarded to N.P., and BBS/E/ER/230001C (Cellular Genomics (CELLGEN): Cell-type specific transcriptional changes during development and upon stress). We thank Earlham Institute National Bioscience Research Infrastructure (NBRI) in Transformative Genomics (BBS/E/ER/23NB0006) for access to Illumina sequencing.

## Methods

### Design and Construction of the Lentiviral Barcoded Vector EB04119 and Bacterial Culture Preparation for High-Throughput Extraction

We used the EB04119 plasmid that is a modified version of pEGZ2^12,13^ (Figure 1), made Golden Gate cloning compatible and containing a DNA barcode (total size: 10,971 bp). The vector pEGZ2 was domesticated to remove an internal BsmBI restriction site in the hygromycin cassette through a synonymous mutation in valine (GTC > GTT) in the CDS using Gibson assembly. An mScarlet dropout cassette (amplified from pMYT083 addgene #180736) was introduced downstream of Woodchuck Hepatitis Virus (WHV) Posttranscriptional Regulatory Element (WPRE) with outward-facing BsmBI recognition sites. Fusion sites (overhangs) were selected from EMMA (Extensible Mammalian Modular Assembly Toolkit) ^17^, specifically position 17 (GCTC-GCCA). Using this domesticated vector containing the mScarlet cassette, we made additional modifications by PCR-amplifying five fragments for Golden Gate assembly using PaqCI (a non-cutting enzyme in this context). Overhang sequences were verified using NEBridge Ligase Fidelity (https://ligasefidelity.neb.com). The oligonucleotides sequences used were: FR1_Fw - TTCACCTGCTTTTCACATTTCCCCGAAAAGTGC, FR1_Rv - TTCACCTGCTTTTACTAGCAAAGGCGGGGAGGCGGCCCAAA, FR2_Fw - TTCACCTGCTTTTCTTCTGAGGCGGAAAGAAC, FR2_Rv - TTCACCTGCTTTTTGTGCGCGGAACCCCTATTTGTT, FR3_Fw - TTCACCTGCTTTTATGGAATTAATTCTGCAGTCGAGAC, FR3_Rv - TTCACCTGCTTTTGAAGTCGAGGCTGATCAGCGGG, FR4_Fw - TTCACCTGCTTTTCGGTCTGTGCCTTCTAGTTGCC, FR4_Rv - TTCACCTGCTTTTCCATAGAGCCCACCGCAT, FR5_Fw - TTCACCTGCTTTTTAGTCTACGCTCTGAGACGGAAAGTG, FR5_Rv - TTCACCTGCTTTTACCGTGAGACGTATAAAC.

PCR products were cleaned up (Zymo kit; eluted in 15.5 μL water) and Golden Gate assembly was performed following NEB protocol (https://www.neb.com/protocols/2021/01/11/golden-gate-assembly) with cycling conditions: (37°C, 1 min → 16°C, 1 min) × 30-60 cycles → 37°C, 5 min → 60°C, 5 min. The assembly mix (2.5 μL) was transformed into 35 μL NEB 10-beta competent cells. Positive clones were selected by colony PCR and fully sequenced (labeled as EB04119). Vector EB04119 was used in a subsequent Golden Gate reaction with annealed barcoded oligonucleotides to insert the barcode sequence, generating a validated plasmid which serves as an example of a final barcoded vector. Glycerols stocks (transformed E. coli, strain Subcloning Efficiency DH5 Competent Cells (ThermoFisher) suspended in 40% glycerol) containing the plasmids of interest were stored in FluidX tubes in the −70°C C automatic freezer arktic (SPT Labtech). The glycerol stocks were retrieved from the arktic in a plate, a decapper Aperio 8 Channel Semi-Automatic Decapper was used to delid tubes. Cultures were inoculated into a 96-deep well plate containing 1.5 ml 2-YT of media plus antibiotic per well, using a multichannel pipette. Three wells (A1, D6, and H12) remain uninoculated and serve as negative controls, confirming the absence of cross contamination across wells. Plate breathable seals we used. The 96-deep well plate was grown overnight at 37°C at 200 rpm in an Innova shaker. The next day, plates were centrifuged at 4,000 g for 30 min at RT and supernatant was discarded. Plates can be stored at −20°C or used immediately for extraction.

### CyBio FeliX Robotic Platform Setup and Labware Configuration

The CyBio FeliX Composer software was used to translate the manual CosMCPrep protocol steps into an automated workflow with minor modifications. The Extraction head is mounted on the CyBio FeliX. A 12-well reservoir plate Reagent Trough (StarLab 12 Channel Trough E2310-1200) was prepared containing the kit CosMCPrep Plasmid Purification Kit A37064 reagents (RE1, L2, N3), fresh 80% ethanol, freshly prepared isopropanol with beads, and elution buffer (EB) The plate is filled according to the number of samples and added to position 9 in the deck (Figure S1). Labelled labware is positioned as in the initial deck layout: The sample plate (containing bacterial pellets) is positioned onto the shaker in position 1, 8-channel adaptor in position 3, Gripper in position 6, Elution plate (catalogue: E1403-5200), 96-Well PCR Plate, Skirted, Low Profile) in position 8, Binding plate 0.5 ml PlateOne® 96 Deepwell Plate, Round Wells with Round Bottoms (Catalogue: S1896-5000) in position 9, AlpAqua magnet in position 10, AxyVoir reservoir trough for waste in position 11. Tip stand OL3317-11-140, Tip stand OL3317-11-105. Note: references can change from suppliers. Labware needed per 96-well extraction: all labware used per 96-well extraction run is listed below. CyBio FeliX 96/1000 µl tips (144 tips; AnalytikJena, cat. nos. OL3811-25-539-N, OL3810-25-871, and OL3810-25-874) and CyBio RoboTipTray 96/250 µl deep well tips (96 tips; AnalytikJena, cat. no. OL3810-26-661) were used for all liquid handling steps. A 96 deep well plate (Azenta, cat. no. 4ti-0132) was used for bacterial culture processing, alongside a 0.5 ml 96 round well plate (StarLab, cat. no. S1896-5000) as the binding plate and a 96 shallow well plate (Azenta, cat. no. 4ti-0116/0117) as the elution plate. A 12-channel reagent trough (StarLab, cat. no. E2310-1200) was used for dispensing buffers.

### Detailed Automated Plasmid Purification Workflow on the CyBio FeliX Platform

The protocol is run from the CHOICE Felix Composer’ software, by selecting the protocol file. The required drive calibration is executed as guided by the protocol. 250 µL tips are used to add 100µL of RE1 into each well using the 8-channel adaptor (used in all following steps). The plate is shaken for 4 minutes at 1000 rpm. Next, 100µL of L2 are added to each well and incubated at room temperature for 5 min. Then, 120µL of N3 are added and the plate is shaken for 5 minutes at 600 rpm. The protocol pauses and requires a spin step of 30 minutes at 4,000 g at 4 C. The plate is placed back in the shaker module, and the protocol is allowed to continue. While the plate is centrifuging, the bead plate is prepared by adding 132µL from the 12 well reservoir, well 5 and 6, into the binding plate in position 9. Once the sample plate is back on the deck 175µL of supernatant is transferred into the bead-containing binding plate. The binding plate is then moved onto the magnetic rack and allowed to settle for 8 minutes. The supernatant is then discarded into the waste position. The next steps involve 3 washes utilizing 100µL of 80% ethanol while the plate remains in the magnetic rack position. After the last step, the binding plate is moved onto the heater shaker and incubated at 37°C degrees for 5 min. The protocol incorporates a checkpoint where visual inspection is needed to confirm ethanol evaporation. If necessary, the plate can be further incubated at 30°C for 5–10 minutes until complete evaporation of residual ethanol is achieved. The plate is placed back on the magnet module and moved to position 8. A customizable volume of elution buffer is added, and the plate is left incubated at 37°C degrees for 5 min on the heater shaker. After incubation, the plate is placed back on the magnetic rack and allowed to settle for 2 min. The eluate is then transferred to the final eluate plate in position 9. The protocol is then completed and ready for further quantification.

### Plasmid DNA quantification, normalisation and E-Gel™ 96Agarose Gels

Plasmid DNA was quantified using an 8-channel NanoDrop and the Quant-iT™ dsDNA Assay Kit Broad Range (BR) (ThermoFisher, Q33130), with fluorescence measured on a CLARIOstar® Plus Microplate Reader (BMG Labtech). Reactions were miniaturised to a 50 µL volume (buffer and reagent at 1:200) with 1–2 µL of DNA sample, set up in a black 384-well plate (Azenta, 4ti-0264). Standard curve analysis was performed in Excel by plotting standard readings, fitting a linear trendline, and extracting the equation (y = mx + b) and R² value. Sample concentrations (ng/µL) were calculated using the inverse equation: x = [(y − b)/m] × dilution factor. DNA samples were subsequently normalised to 10 ng/µL in a 20 µL final volume using an Opentrons Flex protocol adapted from the Opentrons library (https://library.opentrons.com/p/dna-normalization-flex-pipettes). The protocol uses 50 and 1000 µL single-channel pipettes to dispense water followed by DNA, with volumes calculated automatically from a CSV input file containing sample concentrations (ng/µL). Samples were visualised on E-Gel™ 96 Agarose Gels (ThermoFisher) using a Hamilton MicroLab integrated with an E-Base™ Electrophoresis Device (ThermoFisher) (Figure S3). Images were edited using E-Editor software.

### Validation of the Barcoded Plasmid Library by High-Throughput Amplicon Sequencing and analysis

Oligonucleotides were designed to amplify a 172 bp region encompassing the barcode, while incorporating a unique 9 bp tag to enable barcode identification and tracking, in preparation for Illumina library sequencing and barcode complexity analysis. A total of 48 unique forward oligonucleotides alongside 11 reverse oligonucleotides were designed (Table S1), allowing for all PCR products to be uniquely tagged. PCR amplification was performed across the pooled and normalised 96 well DNA plates, giving a total of 528 PCRs, with PCR amplification confirmed via E-Gel DNA electrophoresis. Samples that failed to amplify were subjected to manual PCR repeat. Normalised DNA was diluted to 0.4 ng/µl for use as a template for PCR. Reactions were set up in a final volume of 20 µl using a Viaflo (Integra) as follows: 2 µl DNA (0.4 ng/µl), 2 µl of each oligonucleotide (10 µM), 10 µl DreamTaq Green PCR Master Mix 2X (Thermo Fisher), and 4 µl H₂O (oligonucleotide sequences in Table S1). Cycling conditions: 2 min at 95°C; 30 s at 95°C, 6 s at 74°C (×30 cycles); 3 min at 74°C. PCR products (2 µl) from all 6 × 96-well plates were pooled into a single tube using the Viaflo and purified twice with AMPure XP beads (Beckman Coulter): first at 1.8× ratio (eluted in 40 µl elution buffer), then at 1.2× ratio (eluted in 0.1× TE buffer), with two 70% ethanol washes each time. Concentration and purity were assessed by NanoDrop and Qubit. Cleaned PCR products (200 ng) were used for library construction with the NEBNext Ultra Express DNA Library Kit (3 PCR cycles). Libraries were quantified by qPCR (KAPA Library Quantification Kit, Roche), molarity adjusted using Bioanalyzer (Agilent) data, and sequenced on an Illumina NextSeq 1000 P1 flow cell (300 cycles, XLEAP-SBS) at the NBRI Transformative Genomics facility, Earlham Institute.

The sequencing data was then analysed, for this, the paired-end reads were joined using BBMerge^18^, with over 97% reads merging into single reads. Reads containing potential barcodes must fully match the following structure: index1+const1+[ACGT]*+barcode_pattern+const2+index2, where index1 and index2 are combinatorial index tags (48 forward and 2 reverse, respectively) that together uniquely identify a sample’s well ID, const1 and const2 are constant PCR oligonucleotide sequences; and barcode_pattern is a semi-random sequence used for barcode identification with a design inspired by Bystrykh, L. V et al ^19^. A modified version of the function CellBarcode ::bc_extract ^20^ was utilised, firstly to identify reads of this structure, then, from the candidate reads to extract barcodes fully matching the barcode pattern. Next, a group of barcodes with Levenstein distance of 1 were merged as one. This step largely removes errors introduced by PCR amplification and sequencing^17^. Barcodes each contributing less than 2% of a well’s candidate reads were considered being low frequency and were filtered out. Wells were filtered to remove ones with less than 10,000 candidate reads or with barcodes appearing in more than one well. The library’s barcode list consists of all final barcodes from the filtered wells (Supplementary Data). In total we validated 2,255 barcoded plasmids by sequencing on top of being able to see fluorescence after transduction of cell lines, this indicated that the pipeline is functional and constitutes a new automated tool for lineage tracing barcoded DNA libraries preparation.

### Cell lines Transfection

HEK293T cells (National Gene Vector Biorepository, NGVB, IN, USA) were seeded in 12-well plates (Corning) at 70% confluence in DMEM (high glucose, Gibco) supplemented with 10% FCS (Gibco), 1% penicillin-streptomycin (Gibco), and 1% L-glutamine (Gibco). Cells were transfected using GeneJuice (Sigma-Aldrich) with the following vectors: psPAX2 (Addgene, #12259), pCMV-VSVG (Addgene, 8454), and EB04119. Viral supernatants were collected at 48 and 72 hours post-transfection, centrifuged at 6000 g for 5 min, and used immediately for transduction^12^.

## Supporting Information

**Table S1.**
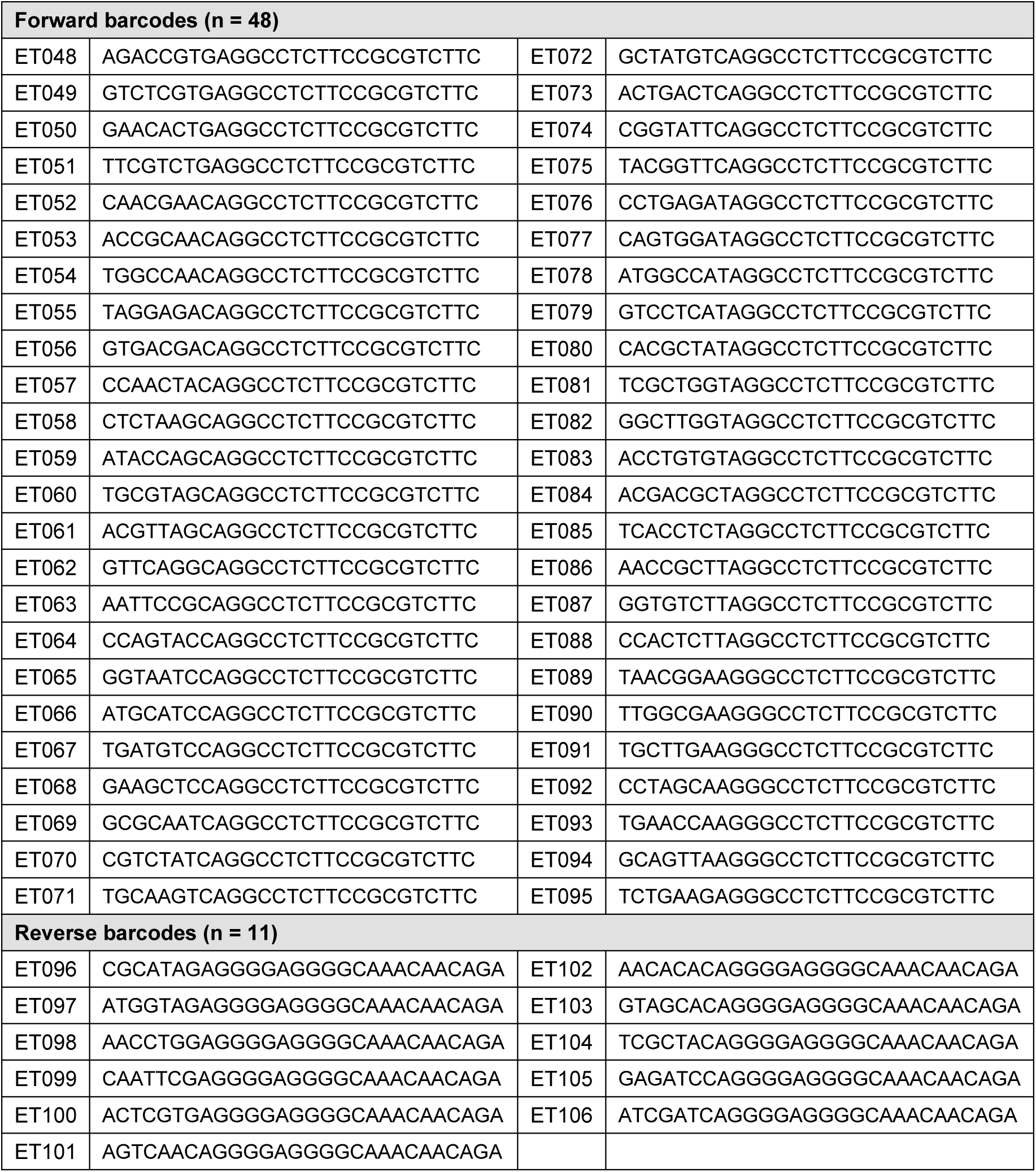
Oligonucleotides utilised for barcode amplification for Illumina sequencing.

**Table S2.** Barcode detection on Plate 1-5 using the bioinformatic pipeline. Number of reads and barcodes processed in each bioinformatic step. The aerosol filtering removed read sequences that appeared only once. Candidate reads were reads match the read structure “index1+const1+[ACGT]*+barcode_pattern+const2+index2”. Raw barcodes were extracted from candidate reads and had a complete match to the barcode pattern. Barcodes were clustered using Levenstein distance before filtering.

| PlateID | Merged reads | Deduplicate d reads | Aerosol filtered reads | Candidate reads | Raw BCs | Clustered BCs | Final BCs |
| --- | --- | --- | --- | --- | --- | --- | --- |
| plate1 | 5,603,546 | 599,983 | 223,077 | 86,822 | 39,753 | 28,040 | 286 |
| plate2 | 14,730,393 | 1,437,738 | 485,657 | 165,665 | 71,460 | 48,800 | 249 |
| Plate3 | 3,621,128 | 439,816 | 166,634 | 64,018 | 29,968 | 20,998 | 133 |
| Plate4 | 8,414,770 | 805,726 | 306,430 | 115,274 | 49,536 | 33,527 | 297 |
| Plate5 | 19,961,803 | 1,678,229 | 548,777 | 196,862 | 88,115 | 61,006 | 296 |

**Table S3.**
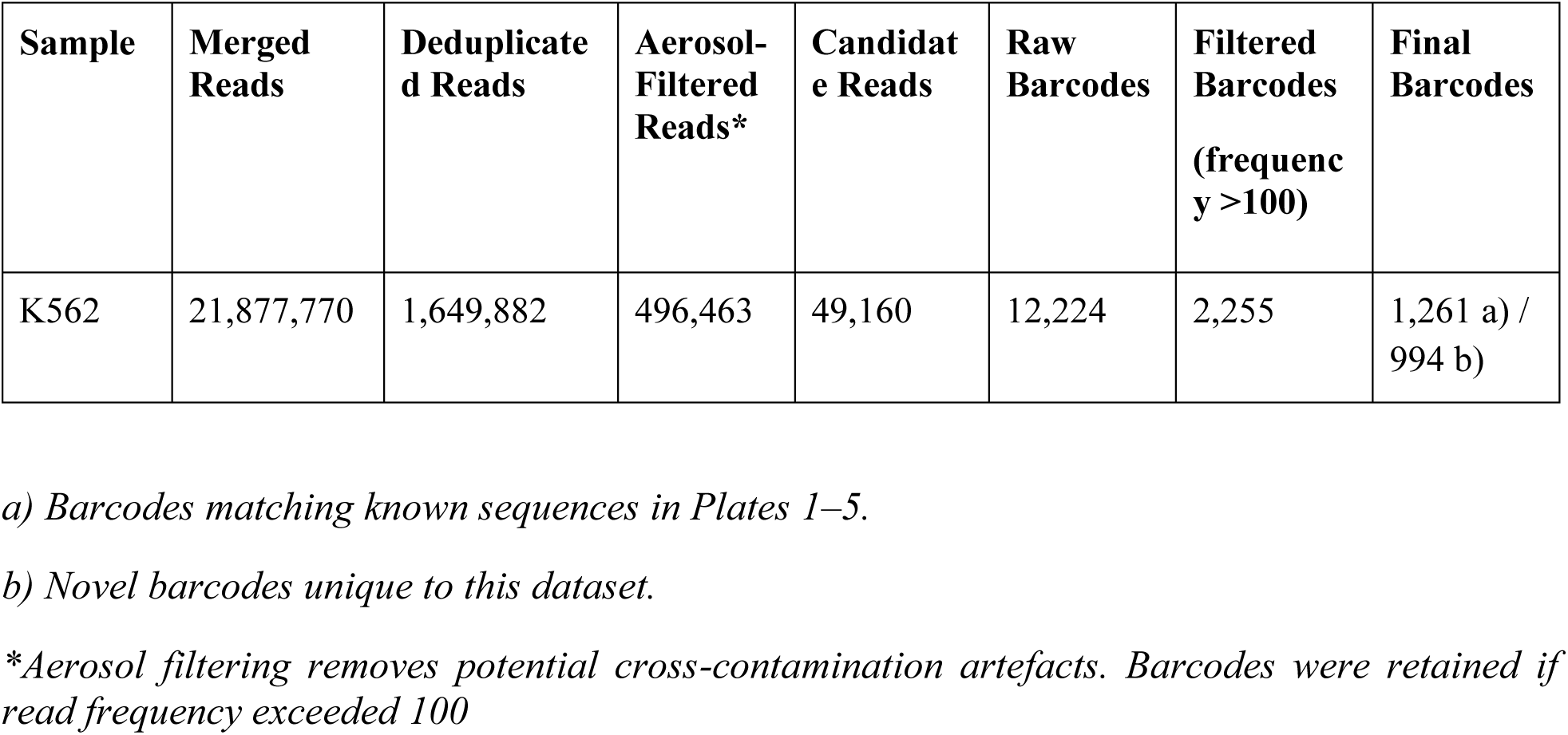
Barcode detection on K562 well using the bioinformatic pipeline. K562 well was processed in the same manner as Table S2 up to the raw BC matching step, except that the maximum edit distance of 3 was allowed. Some of the final barcodes were also found in the final barcode list in Plate 1-5, whereas others were unique to this sample, and had been counted towards the library barcode whitelist.

**Figure S1.**
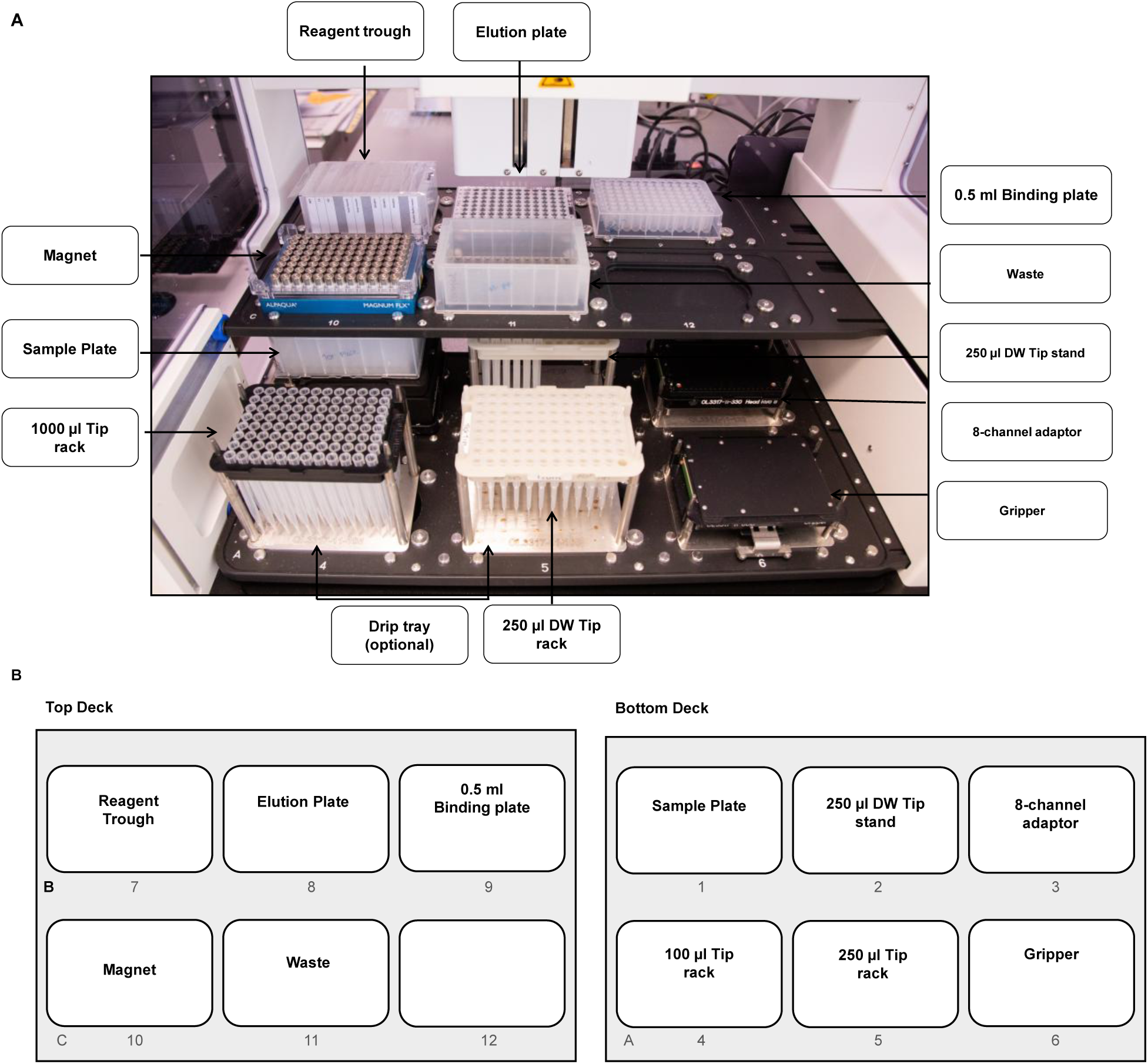
CyBio FeliX robotic platform deck layout for automated plasmid DNA purification. (A) Photograph of the CyBio FeliX liquid handling robot deck configured for high-throughput SPRI-based plasmid DNA purification. Key components are labelled, including the sample plate, reagent trough, elution plate, 0.5 ml binding plate, magnet, waste reservoir, tip racks (1000 µl and 250 µl), 250 µl deep well tip stand, 8-channel adaptor, gripper, and optional drip tray. (B) Schematic representation of the robotic deck layout showing the precise positioning of all components across deck positions A–C (rows) and 1–12 (columns). Sample plate (A1), 250 µl DW tip stand (A2), 8-channel adaptor (A3), 1000 µl tip rack (A4), 250 µl tip rack (A5), gripper (A6), reagent trough (B7), elution plate (B8), 0.5 ml binding plate (B9), magnet (C10), and waste reservoir (C11).

**Figure S2.**
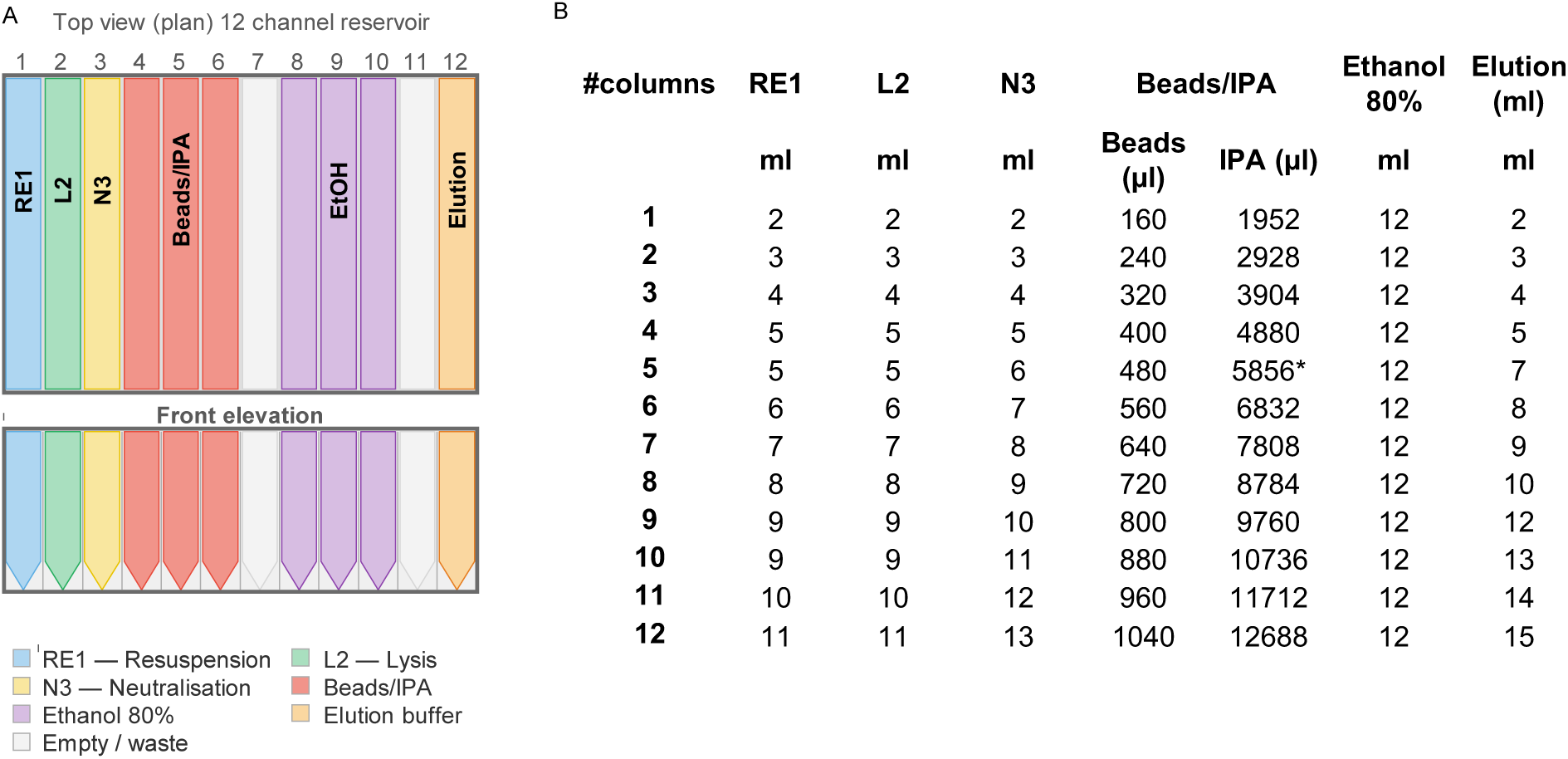
Reagent through layout and volume preparation for the automated SPRI-based plasmid purification protocol. (A) Schematic of the 12-channel reagent trough (StarLab, cat. no. E2310-1200) showing the distribution of CosMCPrep kit reagents (Beckman Coulter) across wells 1–12. Wells are loaded as follows: well 1: resuspension buffer RE1; well 2: lysis buffer L2; well 3: neutralisation buffer N3; wells 4–6: magnetic bead/isopropanol (Beads/IPA) mixture; wells 8–10:80% ethanol (three wash steps); well 12: elution buffer (EB). Highlighted wells (salmon) indicate positions reserved for waste or unused during the run. (B) Reference table indicating the volumes of each reagent to be added to the trough depending on the number of samples processed per run.

**Figure S3.**
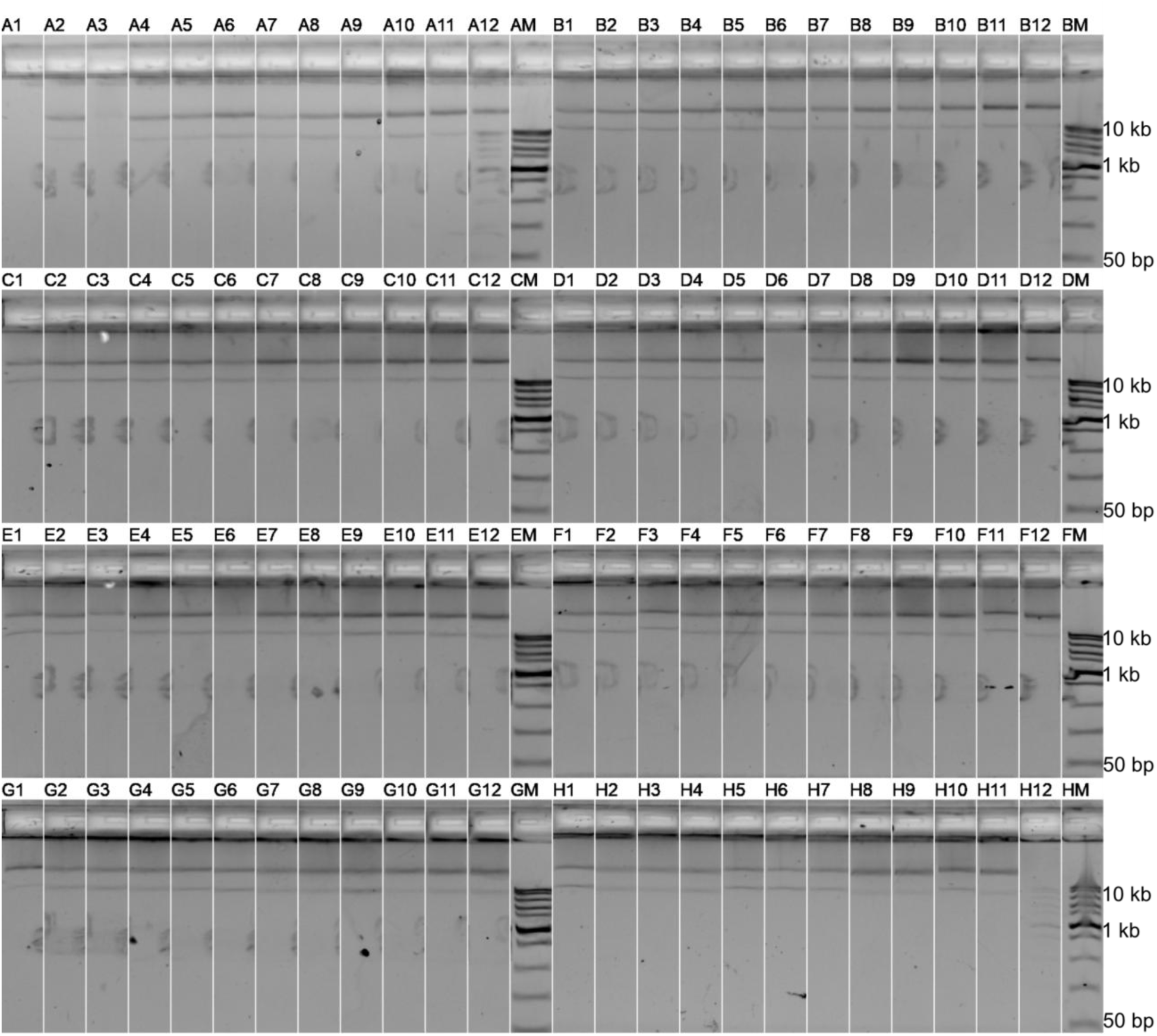
E-Gel™ 96 Agarose Gel loaded with 96 plasmid DNA minipreparation samples, arranged in rows A–H (columns 1–12), with the Fast DNA Ladder (NEB #N3238) (M) included in each row for size reference. The barcoded vectors are 9.9 kb long.

## References

1. Nora, L. C., Westmann, C. A., Martins-Santana, L., Fátima Alves, L., Oliveira Monteiro, L. M., Guazzaroni, M. E. & Silva-Rocha, R. The art of vector engineering: towards the construction of next-generation genetic tools. Microbial Biotechnology 12 (2019). 10.1111/1751-7915.13318

2. Chao, R., Mishra, S., Si, T. & Zhao, H. Engineering biological systems using automated biofoundries. Metabolic engineering 42 (2017). 10.1016/j.ymben.2017.06.003

3. Yang, C.-C., Deshpande, A. J., Jackson, M., Adams, P. D., Yin, J.-A., Wu, Y., Knuff, C. J., Ghias, A., Beketova, A. & Huang, C.-T. Automation workflow for high-throughput arrayed plasmid DNA preparation and quantification. bioRxiv (2025). 10.64898/2025.12.13.694144

4. Vegh, P., Donovan, S., Rosser, S., Stracquadanio, G. & Fragkoudis, R. Biofoundry-Scale DNA Assembly Validation Using Cost-Effective High-Throughput Long-Read Sequencing. ACS Synthetic Biology 13 (February 8, 2024). 10.1021/acssynbio.3c00589

5. Cohen, M., Randolph, D. E., Lozano, M. E., Anderson, P. W., Crissman, J., Triana, F. J. & Cujec, T. A fully automated high-throughput plasmid purification workstation for the generation of mammalian cell expression-quality DNA. SLAS technology 27 (2022 Aug). 10.1016/j.slast.2022.01.005

6. Holowko, M. B., Frow, E. K., Reid, J. C., Rourke, M. & Vickers, C. E. Building a biofoundry. Synthetic Biology 6 (2021/02/17). 10.1093/synbio/ysaa026

7. Hillson, N. et al. Building a global alliance of biofoundries. Nat Commun 10, 2040 (2019). 10.1038/s41467-019-10079-2

8. Yu, C. Y., Ang, G. Y., Mat Isa, N. & Chan, K.-G. Frontiers in biofoundry: opportunities and challenges. Frontiers in Synthetic Biology 3 (2025/10/03). 10.3389/fsybi.2025.1630026

9. Heo, Y. B., Park, J. S. & Woo, H. M. Architectures of emerging biofoundry platforms for synthetic biology. Current Opinion in Biotechnology 96 (2025/12/01). 10.1016/j.copbio.2025.103379

10. Billeci, K., Suh, C., Di Ioia, T., Singh, L., Abraham, R., Baldwin, A. & Monteclaro, S. Implementation of an Automated High-Throughput Plasmid DNA Production Pipeline. Journal of laboratory automation 21 (2016 Dec). 10.1177/2211068216630547

11. Kim, H., Hillson, N. J., Cho, B.-K., Sung, B. H., Lee, D.-H., Kim, D.-M., Oh, M.-K., Chang, M. W., Jin, Y.-S., Rosser, S. J., Vegh, P., Fragkoudis, R., Le Feuvre, R., Scrutton, N. S., Storch, M., Seong, W., Freemont, P. S. & Lee, S.-G. Abstraction hierarchy to define biofoundry workflows and operations for interoperable synthetic biology research and applications. Nat. Commun. 16 (2025). 10.1038/s41467-025-61263-6

12. Wojtowicz, E. E., Mistry, J. J., Uzun, V., Hellmich, C., Scoones, A., Chin, D. W., Kettyle, L. M., Grasso, F., Lord, A. M., Wright, D. J., Etherington, G. J., Woll, P. S., Belderbos, M. E., Bowles, K. M., Nerlov, C., Haerty, W., Bystrykh, L. V., Jacobsen, S. E. W., Rushworth, S. A. & Macaulay, I. C. Panhematopoietic RNA barcoding enables kinetic measurements of nucleate and anucleate lineages and the activation of myeloid clones following acute platelet depletion. Genome biology 24 (06/27/2023). 10.1186/s13059-023-02976-z

13. 13. Belderbos, M. E., Koster, T., Ausema, B., Jacobs, S., Sowdagar, S., Zwart, E., de Bont, E., de Haan, G. & Bystrykh, L. Clonal selection and asymmetric distribution of human leukemia in murine xenografts revealed by cellular barcoding. Blood 129 (2017/06/15). 10.1182/blood-2016-12-758250

14. Oberacker, P., Stepper, P., Bond, D. M., Höhn, S., Focken, J., Meyer, V., Schelle, L., Sugrue, V. J., Jeunen, G.-J., Moser, T., Hore, S. R., von Meyenn, F., Hipp, K., Hore, T. A. & Jurkowski, T. P. Bio-On-Magnetic-Beads (BOMB): Open platform for high-throughput nucleic acid extraction and manipulation. PLoS biology 17 (01/10/2019). 10.1371/journal.pbio.3000107

15. Suckling, L., McFarlane, C., Sawyer, C., Chambers, S. P., Kitney, R. I., McClymont, D. W. & Freemont, P. S. Miniaturisation of high-throughput plasmid DNA library preparation for next-generation sequencing using multifactorial optimisation. Synth Syst Biotechnol 4, 57–66 (2019). 10.1016/j.synbio.2019.01.002

16. Whitehead, E. Automated Planning Enables Complex Protocols on Liquid-Handling Robots. ACS Synth Biol 7, 922–932 (2018). 10.1021/acssynbio.8b00021

17. Martella, A., Matjusaitis, M., Auxillos, J., Pollard, S. M. & Cai, Y. EMMA: An Extensible Mammalian Modular Assembly Toolkit for the Rapid Design and Production of Diverse Expression Vectors. ACS Synth Biol 6, 1380–1392 (2017). 10.1021/acssynbio.7b00016

18. Bushnell, B., Rood, J. & Singer, E. BBMerge – Accurate paired shotgun read merging via overlap. Plos One 12(10) (2017). 10.1371/journal.pone.0185056

19. Bystrykh, L. V. Generalized DNA Barcode Design Based on Hamming Codes. Plos One 7(5) (2012).

20. Sun, W., Perkins, M., Huyghe, M., Faraldo, M. M., Fre, S., Perié, L. & Lyne, A.-M. Extracting, filtering and simulating cellular barcodes using CellBarcode tools. Nat Comput Sci 4, 128–143 (2024). 10.1038/s43588-024-00595-7

